# Semisynthetic Simulation for Microbiome Data Analysis

**DOI:** 10.1101/2024.10.14.618211

**Authors:** Kris Sankaran, Saritha Kodikara, Jingyi Jessica Li, Kim-Anh Lê Cao

## Abstract

High-throughput sequencing data lie at the heart of modern microbiome research. Effective analysis of these data requires careful preprocessing, modeling, and interpretation to detect subtle signals and avoid spurious associations. In this review, we discuss how simulation can serve as a sandbox to test candidate approaches, creating a setting that mimics real data while providing ground truth. This is particularly valuable for power analysis, methods benchmarking, and reliability analysis. We explain the probability, multivariate analysis, and regression concepts behind modern simulators and how different implementations make trade-offs between generality, faithfulness, and controllability. Recognizing that all simulators only approximate reality, we review methods to evaluate how accurately they reflect key properties. We also present case studies demonstrating the value of simulation in differential abundance testing, dimensionality reduction, network analysis, and data integration. Code for these examples is available in an online tutorial (https://go.wisc.edu/8994yz) that can be easily adapted to new problem settings.

## 1 Introduction

Microbial communities play a central role in human and ecological health. Advances in sequencing technology and statistical methods have made it possible to characterize these communities at unprecedented levels of precision [37, 31, 82]. However, statistical methods have to contend with microbiome-specific challenges, like sparse read coverage, large library size variations, uncertainties in the resolved taxa, and overdispersion [71, 104, 42]. These difficulties intensify as scientific goals grow more complex. This is especially evident as microbiome data analysis shifts away from descriptive studies towards modelling that often requires experimental designs with multiple batches, longitudinal sampling, or complementary assays [29, 19, 51]. In this context, effective use of statistical methods hinges on many steps: framing precise questions, preparing suitable data, applying appropriate methods, and delivering accurate interpretations.

Given the uncertainties present in real data, guiding analysis using simulations where the ground truth is known can be tremendously helpful. Simulation has a long history in shaping microbiome data analysis. For example, McMurdie and Holmes [71] applied rarefaction to datasets simulated from a negative binomial model to clarify its impact on downstream inferences. Similarly, Kodikara et al. [52] employed an autoregressive model to assess method performance for longitudinal studies. In fact, most methodological research requires benchmarking on simulated data. Despite these strong precedents, the field is only beginning to formalize, evaluate, and share reusable simulators. In particular, recent advances have proposed simulators that leverage existing *template data* – real experimental data whose patterns the simulator should mimic – thus simultaneously reducing development time while improving fidelity. In the remainder of this review, words in italic font are listed in our glossary of terms, Table 1.

**Table 1:** A glossary of simulation terms.

| Term | Definition |
| --- | --- |
| Covariate | A biological or experimental characteristic of a sample that is thought to impact community profiles and is used to train the simulator. |
| Copula | A statistical approach for modeling correlation in non-Gaussian data, often used to model associations between taxa in a community. |
| De Novo Simulator | A simulator whose mechanism is defined without reference to external, template data. |
| Factor Model | A model that induces correlation across features through latent variables. Like copulas, these are often used to induce associations between taxa. |
| Fit-for-Purpose | Approaches that assess simulation quality by referring to small subsets of taxa. |
| Global Evaluation | Approaches that assess simulation quality by referring to the entire community (all taxa). |
| Marginal Distribution | A probability model that reflects abundance patterns for a specific taxon. |
| Multivariate Model | A probability model that reflects abundance patterns across the entire community (all taxa). |
| Outcome-Specific | Evaluation criteria that target the simulator’s downstream application. |
| Reliability Analysis | The use of simulated data to calibrate interpretations of real data analysis. |
| Semisynthetic Data | The output from a simulator that has been designed to mimic external, template data. |
| Template Data | Previously gathered experimental data that can be used to train a simulator. |

This review introduces readers to emerging trends in simulation, walks through example use cases, and distills best practices. While research on simulation methods often emphasizes realism of the simulated data, their applications are typically only discussed at a high level. Here, we instead explore in-depth applications of simulated data. We first review existing packages and highlight their potential pitfalls (see Table 2). We then illustrate how researchers can get the most out of their microbiome data through simulation for various analytical tasks, ranging from effective experimental designs to data analysis strategies. In particular, this review focuses on three use cases: Power analysis, methods benchmarking, and reliability analysis.

**Table 2:** An overview of packages that are available for microbiome community simulation. Models differ according to mechanisms they use to estimate community structure and the scope of applicability to alternative experimental designs. For applications to new environments or designs, the evaluation techniques described in the section ‘Evaluation’ can be used to compare candidates and gauge simulator fidelity.

| Package Name | Probability Mechanism | Experimental Designs | Notes |
| --- | --- | --- | --- |
| CAMISIM [26] | Models taxon abundances with a log-normal distribution and does not account for featurewise correlation. | Supports separate modes for case-control and time series designs, including simulation of technical replicates. | Used in the CAMI benchmarking challenge [92, 72] and can output read-level data (Illumina, PacBio, or ONT). |
| DeepMicroGen [10] | Combines recurrent network and GAN architectures to model multivariate changes over time. | Supports longitudinal designs with regular spacing. | Parameters are difficult to interpret, limiting potential for control. |
| MB-GAN [88] | Trains a GAN to match a template dataset, capturing both marginal and multivariate structures. | Mimics the design of the template data, but cannot be adapted to alternative designs. | Primarily for Centered Log Ratio (CLR) transformed data but applicable to small sample sizes. |
| metaSparSim [79] | Uses a hierarchical model to sample biological composition (Gamma distribution) and technical variability (multivariate hypergeometric distribution). | Does not include covariates during parameter estimation but allows changes post-estimation to reflect experimental perturbations. | Preserves sparsity and mean-variance profiles for individual taxa, with cross-taxa correlation quality depending on Gamma parameters. |
| miaSim [27] | Simulates population dynamics based on various interaction models, including the generalized Lotka-Volterra model. | Supports time series designs with potential environmental interventions. | Accompanied by an interactive Shiny application but cannot be fitted to a template dataset. |
| microbiomeDASim [106] | Draws community profiles from a zero-inflated, truncated multivariate normal distribution. | Designed for longitudinal experiments with user-specified trends (e.g., quadratic and “hockey-stick”). | Only applicable to normalized abundance tables. |
| MIDASim [38] | Estimates marginals from empirical or generalized gamma models, correlating presence-absence patterns and abundances via a copula. | Supports case-control designs with varying sample sizes, sequencing depths, and taxa presence and abundance. | Scalable for both estimation and simulation, with support for template inputs and post-estimation modifications. |
| scDesign3 [97] | Applies GAMLSS to each feature, linking them with a Gaussian or Vine copula. | Can include covariates during estimation, enabling simulation of arbitrary designs. | Originally developed for single-cell applications, with similar models used for microbiome data [85, 110, 43]. |
| SparseDOSSA 2 [68] | Simulates features from a zero-inflated log-normal distribution, linking them with a Gaussian copula with a regularized precision matrix. | Does not incorporate covariates during estimation, but allows post-estimation changes that use them. | Previously validated on human gut and vaginal microbiome data. |
| ZINB-WaVE [87] | Simulates data from a zero-inflated negative binomial model, incorporating known covariates and low-rank latent variation. | Can include covariates during estimation, enabling simulation of arbitrary designs. | Originally developed for single-cell applications, with similar models used for microbiome data [85, 110, 43]. |

**Power analysis** is an important step in experimental design that informs the sample size required to detect different effect sizes, thereby enabling studies to be done with minimal resources, and without compromising scientific integrity and rigor [112]. Several power calculators have been proposed for microbiome studies [50, 23, 83, 11]. However, these tools lack the flexibility of full simulators, which can be adapted to problem- and data-specific contexts. Most power calculators are limited to case-control designs and, without access to template data, they may make questionable distributional assumptions. In contrast, researchers using simulators can ensure their models reflect important properties of template data and can compare various design choices, like class imbalance and sampling times, that go beyond sample size alone.

**Benchmarking** is essential for identifying the most suitable method for a specific experimental design and data type [25]. Simulation is useful to benchmark methods when ground truth is scarce. For example, in batch effect integration, it is important to preserve biological effects, but these can be difficult to pinpoint without simulations that provide some ground truth (illustrated in the section ‘Batch effect correction’). Further, benchmarking new methods on various plausible datasets provides more insight than simply identifying the best performer in a single simulation scenario. To this end, we can train a simulator to emulate real data and then alter it to reflect multiple hypothetical scenarios. Even when ground truth exists, simulations help gauge robustness to dataset perturbations (e.g., reducing the true effect sizes), and testing many simulated scenarios is often easier than collecting multiple real datasets. Formal simulation techniques provide a more automatic approach to benchmarking and allow researchers to focus on data analysis and method selection, rather than spending time programming simulators from scratch.

**Reliability analysis** can be enabled through simulation. Many modern data analysis workflows lack sufficient theory to guide practice, and simulations offer valuable sanity checks. For example, simulation studies have found that sparse canonical correlation analysis (sCCA) can result in high false discovery rates [33, 39]. Although these studies were motivated by neuroscience, sCCA is also widely used in microbiome data integration [7, 90]. In a similar spirit, the section ‘Omics data integration’ highlights how data integration can introduce unexpected artifacts into dimensionality reduction visualizations. Hence, simulations help prevent misinterpretations that could misdirect research priorities.

This review makes the following contributions:

1. **Overview of simulation workflows**: We describe the main ingredients of modern simulation algorithms and associated assumptions. This background ensures that methods are not treated as black boxes and guides their effective use.
2. **Evaluation criteria**: We outline how to assess whether simulated data match the properties of previously observed template data, and how simulated data can inform methodological improvements.
3. **Case studies**: We present realistic case studies demonstrating how simulation can support power analysis, benchmark competing methods, and guarantee reliable conclusions.

This review is accompanied by an online tutorial (https://go.wisc.edu/8994yz). Each chapter of the tutorial starts with a dataset discussion, walks through the process of designing simulators, and applies the resulting models to address common questions about microbiome experimental design, method benchmarking, and result interpretation.

## 2 Simulation Workflows

Simulators vary widely, and their realism and relevance to downstream tasks is context-dependent. To help navigate this landscape, we first categorize simulators and then outline the workflow for building and refining them.

### De novo and template-based simulators

We first note the distinction between *de novo* and template-based simulators. De novo simulators require users to specify parameters related to experimental design and distributional properties. These parameters cannot be automatically matched to real, template data. For example, in the splatter simulator [109], users can include treatment and batch effects, but these are drawn from the simulator’s internal generation mechanism. In contrast, template-based simulators first estimate the impact of sample-level *covariates* using real data. Though this does require an initial template, it can lead to more realistic synthetic data. These data are often called *semisynthetic*, reflecting the influence of the template. As many public catalogs of 16S, metagenomics, and metabolomics data are now available [78, 74], it has become easier for researchers to access relevant template data for various possible analysis. We caution that different research communities have used different terms to describe the concept of template-based simulation. For example, “semisynthetic” is used in metabolomics [105, 81] and microbiome studies [91, 69], while “reference-based” is common in single-cell genomics [62, 12]. Although de novo and template-based simulators differ in their requirements, similar factors guide their application. We will review these workflow considerations next.

### Formulation and application

In the formulation phase (Figure 1A), a simulation model is created by adjusting parameters until the generated data pass a series of evaluation checks. In the application phase (Figure 1B), several altered versions of the simulator can be defined according to the simulation study’s goals. For example, to characterize false discovery rates, we can introduce synthetic negative control features designed to lack associations with the outcome. For each altered version of the simulator, we can generate multiple datasets, apply candidate analysis strategies, and gather summary statistics quantifying their performance. We provide practical examples of both phases in the ‘Case Studies’ section.

**Figure 1:**
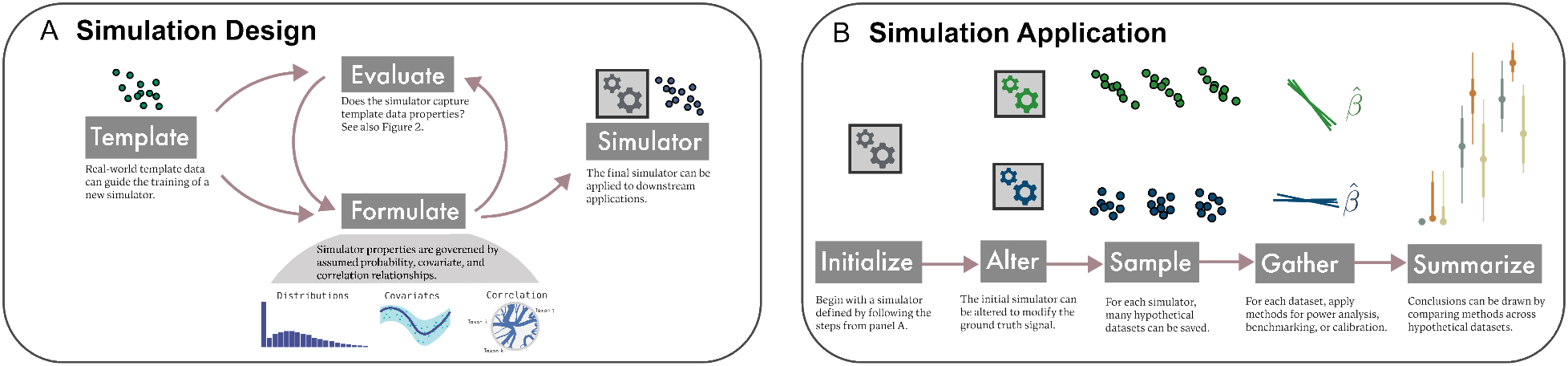
(A) Simulation design is an iterative process, where choices of probability distribution, experimental and biological covariates, and correlation structure can be refined according to evaluation criteria that draw attention to differences between real and template data. See Figure 2 for example criteria. (B) Given a satisfactory simulator, the same workflow applies to power analysis, benchmarking, and reliability analysis. The initial simulator can be modified to reflect hypothetical experimental or biological settings, like changes in the signal effect size or the sample size. Outputs from competing approaches can be gathered and interpreted to guide experimental design and analysis.

The formulation phase relies on concepts from probability, statistical estimation, regression, and multi-variate analysis. Probabilistic models enable sampling of new data with appropriate distributional properties, *multivariate models* induce plausible associations across multiple taxa, and regression methods ensure that simulations reflect biological or experimental influences.

### 2.1 Probability distributions

The choice of probability distribution dictates many properties of the generated data. Distributions must be carefully selected, as properties that might be easy to manipulate in one distributional family might be difficult to modify in another. For example, sparsity can be more easily modified in a zero-inflated vs. ordinary negative binomial distributions. We first review univariate, then multivariate distributions.

#### Univariate distributions for simulation

For a specific taxon, we need to decide whether to simulate counts, proportions, or real numbers. Count distributions can be used to simulate amplicon sequence variant or metagenomics data before having applied any transformations. Common count distributions include the Poisson, negative binomial, and hypergeometric distributions. The Poisson distribution arises when counting events that, though individually unlikely, become common due to repeated opportunities for them to occur. For example, a stretch of DNA is unlikely to align to any given read, but given enough reads, we would expect a Poisson number of them to align. However, in real data, this often gives a poor approximation – the variance in observed counts is often higher than the Poisson can capture. Such overdispersion is more appropriately modeled using the negative binomial distribution. Finally, if the data exhibit more zeros than either Poisson or negative binomial distribution allows, then zero-inflation can be employed to introduce additional sparsity, as in zero-inflated negative binomial (ZINB) used in the single-cell RNA-seq simulator ZINB-WaVE [87]. Alternatively, presence-absence can be modeled separately from abundances, as in the simulator MIDASim [38].

When data are transformed, count distributions no longer apply. Depending on the transformation, different distributions can be used to model nonnegative, interval, or arbitrary real values. For nonnegative values, options include the truncated normal, log-normal, or variants of gamma distributions. The truncated and log-normal distributions enforce nonnegativity by either truncating or exponentiating a normal distribution. These distributions are used in microbiomeDASim [106], SparseDossa 2 [68], and CAMISIM [26]. Proportions within the interval [0, 1] can be represented by beta, Dirichlet, or generalized Gamma distribution, which include parameters for mean and concentration along the boundaries. The simulator MIDASim uses this approach to model relative abundance-transformed data. If data have been transformed to include both positive and negative real values, then they can often be modeled with normal or Student’s T distributions, the latter being more appropriate when outliers are present. Further, such transformations can enable generative adversarial network models to perform well, as in simulators DeepMicroGen [10] and MB-GAN [88]. Continuous distributions can also be used as a preliminary sampling step for hierarchical count models. For example, the means of a count model can first be modeled as a Gamma distribution, allowing estimates for rare taxa to borrow strength from more abundant taxa. This hierarchical approach is used by metaSparsSim [79], which draws Gamma marginal distributions for individual taxa before sampling from a multivariate hypergeometric distribution for all taxa jointly.

#### Multivariate distributions for simulation

It is important that simulators generate realistic community profiles, capturing relational, multivariate structure and not just univariate, per-taxon abundances. Two common strategies for modeling these associations are (1) learning a multivariate normal covariance in an appropriately transformed space or (2) incorporating shared latent variation across features (taxa). Strategy (1) uses the multivariate normal distribution’s covariance parameter to control the relationship between features. These multivariate normal samples can then be transformed into sequencing-specific data distributions. For example, this approach is adopted by simulators using the Logistic-normal Multinomial distribution [2, 111]. This method induces correlations across features by first sampling from a correlated, multivariate normal distribution. This correlated vector is transformed into a vector of proportions using a log-ratio transformation. Given an overall sequencing depth, simulated reads are then allocated to individual taxa based on these proportions. Alternatively, a correlated multivariate normal distribution can be transformed into a multivariate distribution with known univariate marginals, a process known as *copula modeling* [44, 14]. Copula models can be used even when the marginals are drawn from different distributions. This makes copulas easily adaptable, and they are used by simulators MIDASim [38], SparseDOSSA 2 [68], and single-cell simulators scDesign2 [100] and scDesign3 [97].

In contrast, strategy (2) ties features together by assuming a latent, low-dimensional vector. The shared source of variation induces downstream correlations. This is often accomplished through variations of *factor models*. For example, ZINB-WaVE simulates entries from a model whose parameters vary according to a low-dimensional latent vector representing unobserved sample-level properties. Similarly, Latent Dirichlet Allocation assumes that samples are drawn from a multinomial distribution whose mean depends on a low-dimensional community membership vector [89]. In both univariate and multivariate settings, interpretable parameterizations can support modifications of the simulation mechanism. These modifications are helpful for validating workflows through computational negative or positive controls. For example, the mean parameter of a negative binomial model can be adjusted to reflect stronger or weaker treatment effects. In contrast, flexible machine-learning-based multivariate simulators, like MB-GAN and DeepMicroGen, can be challenging to alter in this way. Even if their simulated data are realistic, it is difficult to control them and enforce desired constraints.

### 2.2 Accounting for experimental and biological factors

How can we model differences across experimental or biological conditions? For example, a taxon’s abundance may change gradually over time or may be a marker of disease status. In this case, histograms of individual taxon abundances may reveal complex patterns, like separate peaks for disease and healthy groups, which cannot be captured by models that assume independent draws from the same probability distribution. To simulate these patterns, we can specify distributional parameters based on sample characteristics. In addition to producing more realistic data, conditioning on sample covariates enables more precise control. For example, simulators that take into account time or disease status can generate data at a denser sequence of timepoints or different sample sizes for healthy vs. disease patients, both of which can guide experimental design and method benchmarking.

Many simulators are tailored to specific experimental designs. For example, MIDASim [38] and Sparse-DOSSA 2 [68] support simulation from case-control designs. In both simulators, some taxa share parameters across case and control groups, representing synthetic negative controls. To create a subset of taxa that differ across groups, representing synthetic positive controls, taxa parameters can be allowed to vary. The greater the difference between parameters, the stronger the true effect. These simulators are particularly useful for power analysis and benchmarking for differential abundance studies. Other simulators have been created specifically for longitudinal designs. For example, microbiomeDASim [106] allows taxonomic abundance to vary over time based on various plausible trends, to mimic brief disruptions or gradual development. Similarly, miaSim [27], CAMISIM [26], and DeepMicroGen [10] are designed to capture longitudinal dynamics.

Though these simulators streamline work with specific experimental designs, it can be useful to generalize and map arbitrary sample-level variables onto distributional properties. This is especially valuable in multifactorial experiments, where multiple biological or experimental characteristics can jointly influence measurements. Such generalization also allows modeling interactions or nesting between variables. For example, treatment and control groups may have divergent temporal trajectories, or treatments might have different effects across cohorts. For these applications, parameters can be linked to sample covariates using regression techniques. For example, in the scDesign3 simulator for single-cell and spatial omics data [97], mean and dispersion parameters are modeled as functions of sample characteristics using generalized additive models of location, scale, and shape (GAMLSS) [86, 98]. This flexibility allows simulated data to vary smoothly over temporal or spatial coordinates. Similarly, in the ZINB-WaVE simulator for single-cell RNA-seq data [87], sample-level covariates can influence the mean, dispersion, and zero-inflation parameters through a linear model. Beyond feature-level properties, sample covariates can modify multivariate relationships. For example, scDesign3 allows different copula covariances to be used within prespecified groups of samples.

## 3 Evaluation

Evaluating synthetic data is a crucial step in data simulation to assess whether the generated synthetic data closely mirror the statistical properties of the original data. Without proper evaluation, synthetic data may not accurately represent the underlying distribution, potentially leading to biased models or incorrect conclusions.

In evaluating synthetic data, three primary types of utility measures can be used: *fit-for-purpose measures*, *global (broad) utility measures*, and *outcome-specific (narrow) utility measures* [15] (see Figure 2). Fit-for-purpose measures provide an initial evaluation of whether the synthetic data appear reasonably close to the real data, and they can be used to improve the simulation approach [15]. In contrast, global utility measures aim to assess the multivariate characteristics of the data. Outcome-specific utility measures aim to quantify similarity in analysis results or specific model parameters between real and synthetic data.

**Figure 2:**
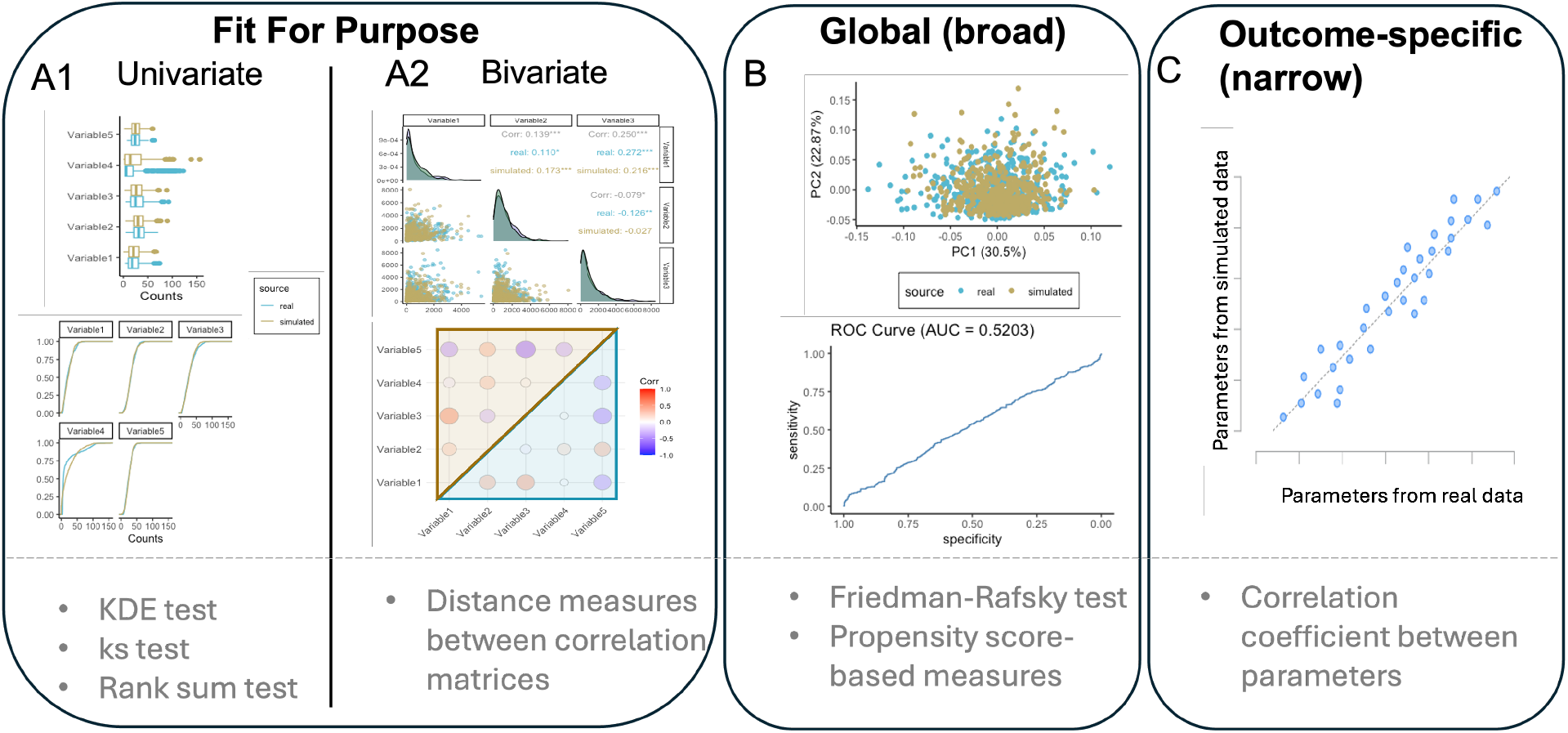
Utility measures for synthetic data, including visual comparison (top) and quantitative measures (bottom). (A) Fit-for-purpose, (B) global (broad), and (C) outcome-specific (narrow) utility measures. Fit-for-purpose and global utility measures both evaluate the similarity between synthetic and real data without considering any analytical objective. In contrast, outcome-specific utility assess the similarity of analysis results or specific model parameters using both real and synthetic data.

Fit-for-purpose measures typically involve checking the univariate and bivariate distributions of observed and synthetic data. We divided these measures into two main types: univariate and bivariate evaluations (Figure 2A1-A2). In the former, we focus on whether the marginal distributions of variables in the real and synthetic data match. In the latter, we focus on whether the pairwise relationships of variables in the synthetic data resemble those in the real data, typically through bivariate distributions and correlations. We further divide evaluation criteria into graphical and quantitative measures. For example, graphical measures include side-by-side box plots or empirical cumulative density plots (univariate setting) and pairwise scatter plots or correlation heat maps (bivariate setting). Quantitative measures include the Kolmogorov–Smirnov (KS) test, the Wilcoxon Rank-Sum test, or independence tests based on kernel density estimation [17] (univariate setting) and the distance measures between correlation matrices (bivariate setting). However, while fit-for-purpose measures offer an initial assessment of synthetic data quality, they do not account for the multivariate nature of the data.

Global utility measures are built upon fit-for-purpose measures to evaluate the complex, multivariate nature of the data. We again divide these into graphical and quantitative measures (Figure 2B). For example, graphical measures include Principal Component Analysis (PCA) plots, which jointly project synthetic and real data into a two-dimensional space, or receiver operating characteristic (ROC) curves, which use binary classification to separate synthetic from real data, to determine if the synthetic and real data can be distinguished. Quantitative measures include the Friedman-Rafsky test [24] or propensity score-based [21] techniques. Global utility measures provide an aggregated similarity between simulated and real data. However, they do not guarantee that specific analyses on real and simulated data will yield similar results, as these measures do not consider a specific analysis goal [15, 49, 20].

Outcome-specific utility measures are designed to assess the simulated data for a particular analysis goal. Since there are multiple analytical approaches, these utility measures can vary significantly. For example, if the focus is on fitting a multiple regression model between sequencing features and a biological condition of interest, then we can compare regression coefficients obtained when fitting the regression to either simulated or real data. In this situation, a scatter plot can serve as a graphical measure, and the correlation value between the parameters estimated from real and simulated data can serve as a quantitative measure (Figure 2C). However, if the objective is to perform a correlation analysis, like those which underlie microbiome network models, then the evaluation should focus on comparing the correlation matrices between the features in the real and the simulated data (for an example of this comparison, see Section ‘Power analysis for multivariate methods’). No single utility measure is universally applicable. Therefore, performing several utility measures based on the specific objective of the simulation will help modify certain aspects of the simulator, such as the selection of distributions.

## 4 Case Studies

### 4.1 Statistical power for univariate differential abundance methods

#### Motivation

Differential abundance analysis is a cornerstone of microbiome research. It is often used to highlight taxa that are associated with disease or that respond to interventions. The research community has developed various testing methods to account for specific characteristics of microbiome data, such as zero-inflation, overdispersion, library size differences, compositionality, and small sample sizes [32, 52, 38]. However, assumptions that are reasonable in one microbiome system (e.g., the gut) may not necessarily translate to others (e.g., marine). Applying a method in an inappropriate context can lead to excess false positives or reduced power. Moreover, unlike classical two-sample tests or linear models, differential abundance methods do not come with closed-form formulas for calculating statistical power. Therefore, semisynthetic simulation can give insights into the properties of differential abundance methods for specific data analysis applications.

We compared three common differential abundance methods commonly used in microbiome studies: DESeq2 [65], limma-voom [57] and ANCOM-BC2 [63]. ANCOM-BC2 was developed specifically for microbiome data, whereas DESeq2 and limma-voom were originally designed for RNA-seq data. The goal of this simulation is to select the most appropriate method for a specific dataset based on statistical power across sample sizes.

#### Data

We analyzed data from the ATLAS study [56]. This study profiled the gut microbiomes of *n* = 1006 healthy adult Europeans. Notably, it discovered microbiome “tipping points,” which are unstable community composition profiles that could “tip” into more stable states. The competing stable states were related to factors like age and body mass index (BMI). We filtered down to the 89 most abundant genera in the 883 subjects with known region membership and BMI within the lean, overweight, or obese categories.

#### Simulation and evaluation

Our simulator follows scDesign3’s modeling assumptions, applying a zero-inflated negative binomial GAMLSS regression where both the mean and dispersion for each taxon can vary as a function of the BMI category. For the top ten most abundant taxa, we found that the boxplot quartiles and the observed shifts across BMI groups were captured well in the simulated data (Figure 3A1). We compared the abundance of all genera in simulated and real data for each BMI group using a kernel density-based two-sample test [16, 18]. At least 70% of genera were not statistically distinguishable (Figure 3A2). As evident by these evaluations, the simulated data were generally similar to the original data. However, for some genera with positively skewed distributions and a high number of outliers, such as *Prevotella melaninogenica et rel*, the simulation failed to capture the high degree of skewness or generate outliers, hence the significant differences between the real and simulated data returned by some of the tests.

**Figure 3:**
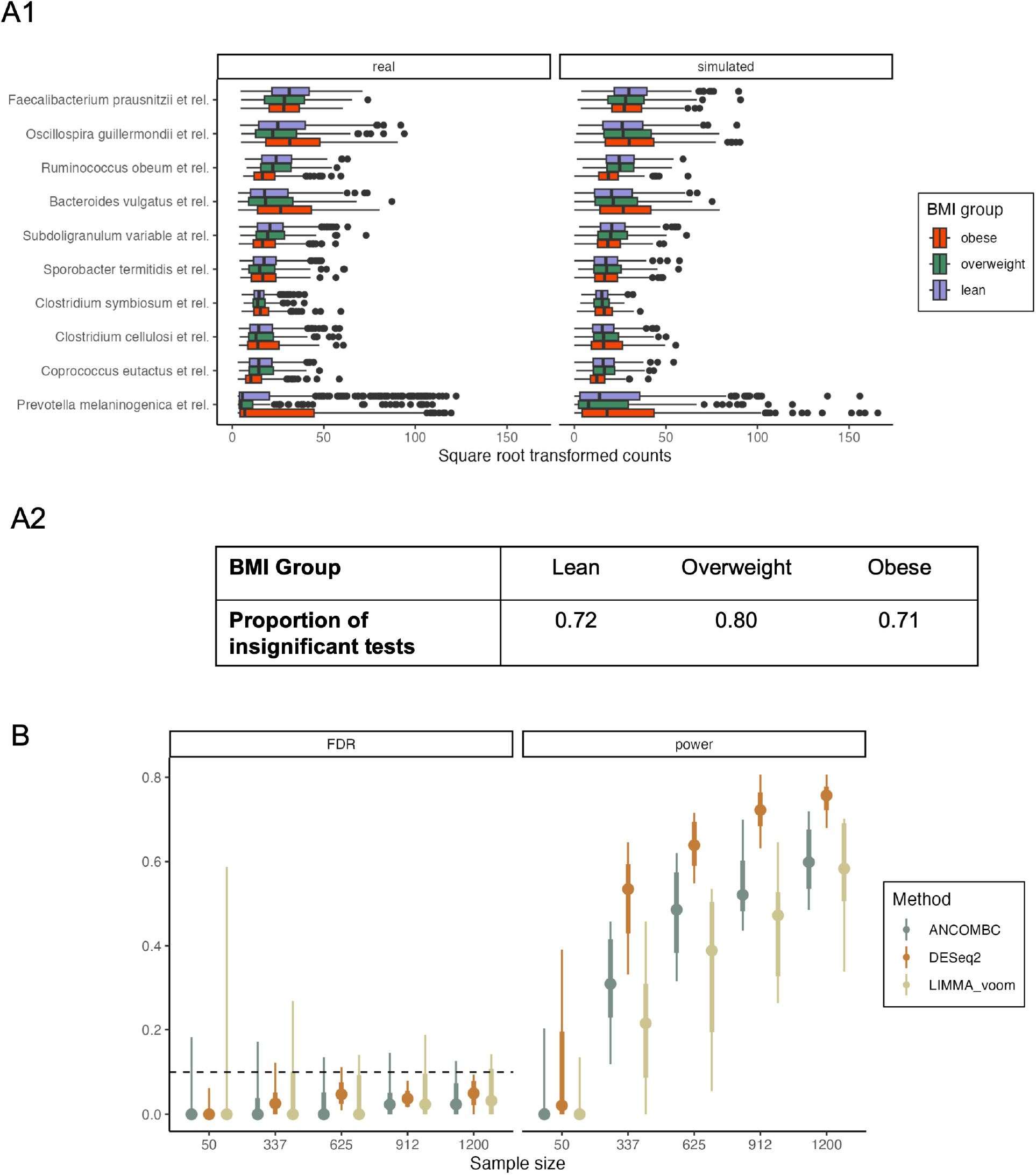
Differential abundance analysis in the ATLAS study. (A1) Comparison between real and synthetic data for the ten most abundant taxa. A zero-inflated negative binomial model achieves satisfactory performance in matching quartiles of the observed data. (A2) Quantitative evaluation between real and synthetic data. Proportion of non-rejected tests from the kernel density-based global two-sample comparison test for each feature across BMI groups in real and simulated data. Insignificant test results indicate that the kernel density estimates for the given feature in each BMI group cannot distinguish between the real and simulated data. (B) Realized power and false discovery rates for the DESeq2, limma-voom and ANCOM-BC2 methods applied to the simulated data. Large samples lead to higher and less variable experimental power. FDR control is maintained on average.

#### Data analysis results

We simulated synthetic negative controls by re-estimating a subset of genera so that the zero-inflated negative binomial means and dispersions did not depend on BMI. These genera were declared not significant from a Wilcoxon Rank-Sum test of association with BMI (*p*-values corrected for multiple testing with 0.1 FDR level). Such weakly associated genera have previously been used to define pseudo-negative controls [76], whereas in our simulator, these genera are genuine negative controls. These simulation design considerations, as well as those for all other case studies, are summarized in Table 4. We then simulated datasets with sample sizes ranging from 50 to 1200, and applied DESeq2, limma-voom and ANCOM-BC2 using an FDR-control level of 0.1. On average, DESeq2 had better power compared to the other two approaches, and all three approaches provided valid FDR control (Figure 3B). The power increased most rapidly from 50 to 337 samples.

**Table 3:** Summary of case study datasets and analysis goals.

| Study name | Number of samples | Number of features | Analysis task | Objective |
| --- | --- | --- | --- | --- |
| ATLAS [56] | 883 | 89 | Differential abundance testing | To select the appropriate differential abundance method for a given experimental design and data type. |
| Type 1 Diabetes [28] | 101 | 427 | Power analysis and multivariate method | To compare complex experimental designs and multivariate analysis approaches. |
| American gut microbiome [70] | 261 | 45 | Benchmarking network inference | To evaluate different network inference approaches under diverse network structures. |
| Anaerobic digestion [6] | 75 | 231 | Batch effect correction | To give insights into the behavior of batch integration methods. |
| Sepsis in ICU patients [36] | 57 | 180 bacteria, 18 fungi, 42 viruses | Omics data integration | To uncover whether different data modalities (bacteria, fungal, virus) should be analyzed together or separately. |

**Table 4:**
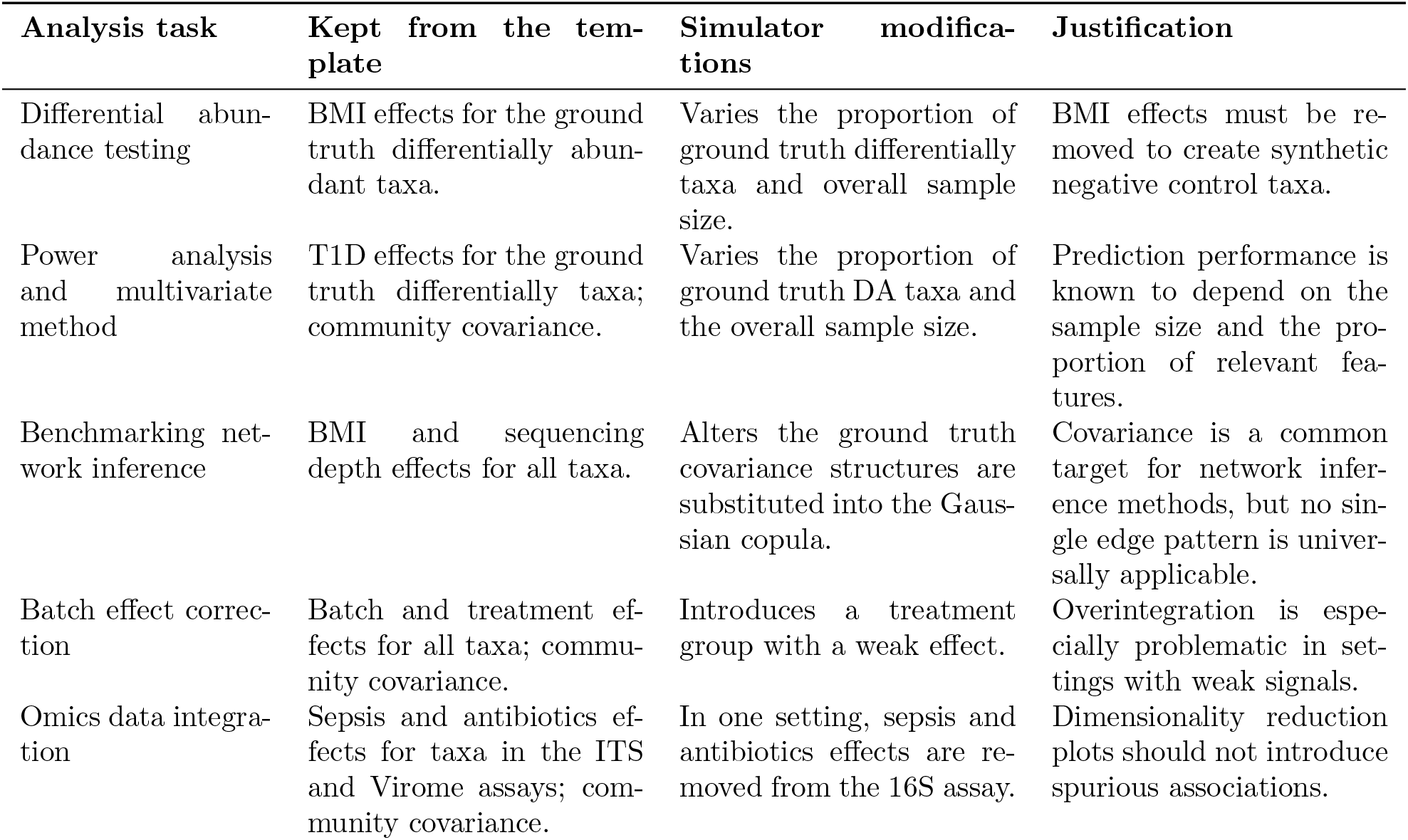
Summary of simulation design considerations for the case studies.

| Analysis task | Kept from the template | Simulator modifications | Justification |
| --- | --- | --- | --- |
| Differential abundance testing | BMI effects for the ground truth differentially abundant taxa. | Varies the proportion of ground truth differentially taxa and overall sample size. | BMI effects must be removed to create synthetic negative control taxa. |
| Power analysis and multivariate method | T1D effects for the ground truth differentially taxa; community covariance. | Varies the proportion of ground truth DA taxa and the overall sample size. | Prediction performance is known to depend on the sample size and the proportion of relevant features. |
| Benchmarking network inference | BMI and sequencing depth effects for all taxa. | Alters the ground truth covariance structures are substituted into the Gaussian copula. | Covariance is a common target for network inference methods, but no single edge pattern is universally applicable. |
| Batch effect correction | Batch and treatment effects for all taxa; community covariance. | Introduces a treatment group with a weak effect. | Overintegration is especially problematic in settings with weak signals. |
| Omics data integration | Sepsis and antibiotics effects for taxa in the ITS and Virome assays; community covariance. | In one setting, sepsis and antibiotics effects are removed from the 16S assay. | Dimensionality reduction plots should not introduce spurious associations. |

#### Summary

Differential abundance testing requires a delicate balance. On the one hand, parametric assumptions, like those for overdispersion or zero inflation, can improve test sensitivity. On the other hand, inappropriate assumptions can lead to invalid results. Generic benchmarking studies can help identify the appropriateness of each method for a given experimental design and data type but can only provide blanket recommendations. In contrast, we showed that simulations let us design “self-service” benchmarks where we can run our own *in silico* experiments to inform a statistical analysis workflow with more problem-specific choices. An additional benefit is that we can ask how certain changes to the data (e.g., increasing the sample size) might affect method performance.

### 4.2 Power analysis for multivariate methods

#### Motivation

Differential analysis methods can draw attention to significant taxa but can overlook important correlation structure within microbiome communities. To shed light on this more global structure, multivariate analysis methods play an essential role [59]. Theoretically characterizing the statistical efficiency of multivariate methods is an active area of research [1, 48], and practical sample size guidance typically relies on simulation [95, 35]. However, effective community-wide simulation is more challenging than what is required for differential abundance analysis, because we must pay close attention to the quality of the estimated correlations. In this case study, we explore how semisynthetic data can inform sample size calculations in an application that uses sparse partial least squares discriminant analysis (SPLS-DA) [58]. SPLS-DA is a classification method that makes use of correlations between input features to ensure stable predictions in small sample size settings. In the process, it computes a dimensionality reduction of the data analogous to PCA, but with the explicit goal of separating classes.

#### Data

We revisited the Type 1 Diabetes (T1D) study from Gavin et al. [28], who identified metaproteomic patterns in the microbiomes of T1D patients from a cohort of *n* = 101 study participants. We filtered down to the 427 proteins present in at least 70% of either the T1D (*n* = 51) or control (*n* = 50) groups. Following the original study’s data preprocessing, we applied a centered log-ratio (CLR) transformation to the measured protein relative abundances and then used these as predictors in an SPLS-DA with T1D status as the outcome. We set the SPLS-DA hyperparameters to 5 PLS dimensions and a selection of 30 predictors. On this dataset, ten repetitions of 5-fold cross validation yields a holdout area under the receiver operating characteristic curve (AUROC) of 0.667 *±* 0.037.

#### Simulation and evaluation

We fitted a Gaussian GAMLSS simulator, allowing means and variances for all proteins to depend on T1D status. Since SPLS-DA models the relationships across all proteins, we first applied a Gaussian copula using the standard sample covariance to attempt to reflect the true correlation structure. Contrary to the last section, we focus here on the quality of the simulated sample correlations rather than the univariate simulation quality (which is covered in our online tutorial section 3.3).

A simple histogram of pairwise correlations between features in simulated data showed greater variability compared to the observed data, suggesting that the simulation could be improved using a regularized covariance estimator. We therefore used the adaptive thresholding covariance matrix estimator of Cai and Liu [5] (higher thresholds apply stronger regularization, while lower thresholds reduce bias). To choose the threshold, we evaluated the correlation quality across a range of candidate values from 0.001 to 0.2 using two metrics: the KS statistic between observed and simulated histograms of pairwise correlations and the Frobenius norm between observed and simulated correlation matrices. The KS statistic formalizes our histogram check, while the Frobenius norm measures the differences between these matrices. We found that the KS statistic was minimized at 0.14, while the Frobenius error was minimized at 0.03. To balance these two metrics, we chose a threshold of 0.1. To further check the simulator, we created pairwise scatter plots (Figure 4A) and heatmaps of real and simulated sample covariance matrices (Figure 4B). The heatmaps show similar blocks of positive or negative correlation. For the pairwise scatter plots, we filtered to a subset of pairs with moderate positive correlations between 0.72 to 0.8 in the real data, as checking all 90K pairs is impossible. The scatter plots of real and simulated data overlapped well. The only notable differences were the streaks of exact zeros in the real data, a reflection of the zeros present before CLR transformation. Overall, the revised simulator with an adaptive covariance estimator seemed sufficient for a power analysis with SPLS-DA.

**Figure 4:**
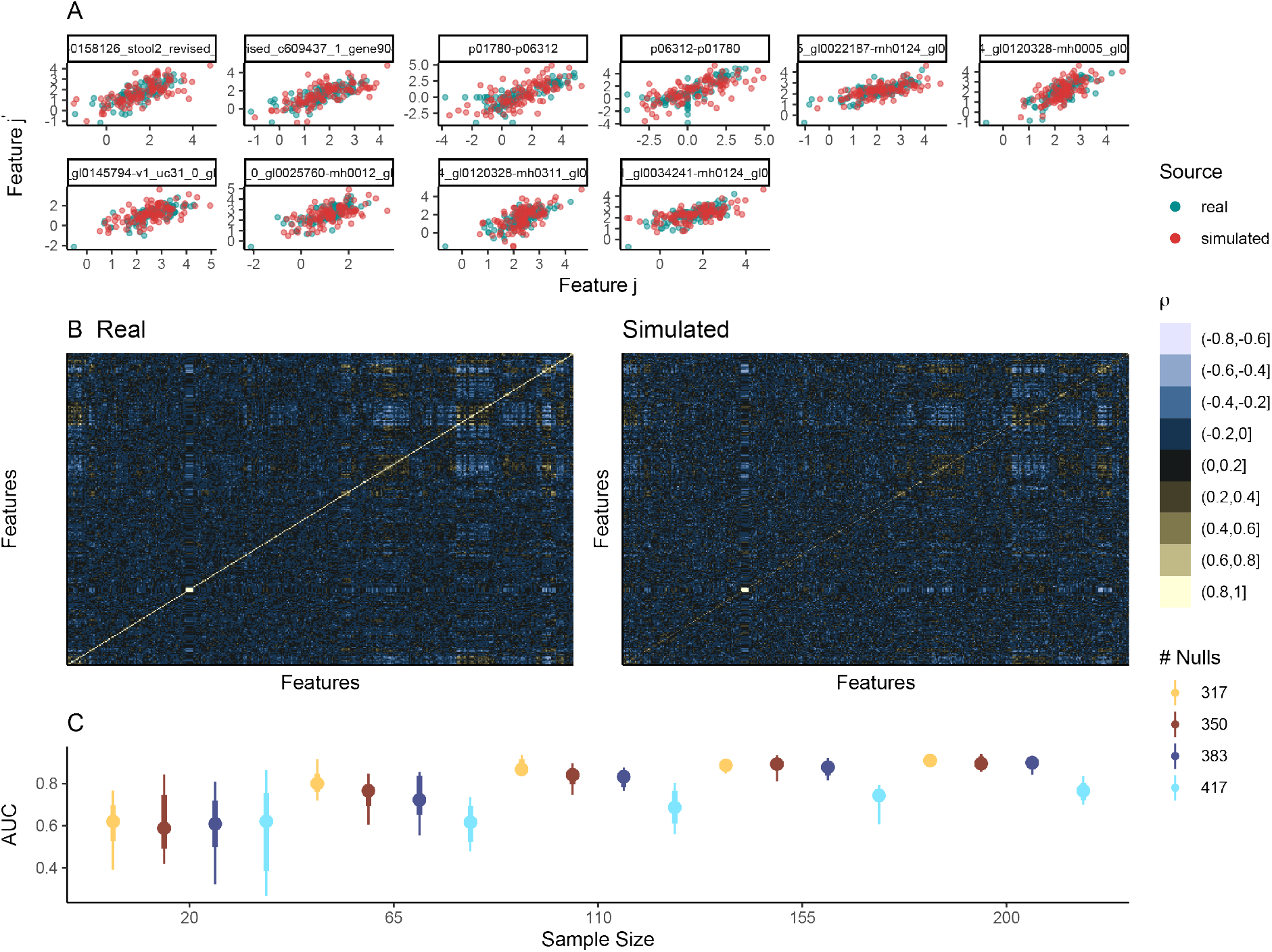
SPLS-DA power analysis for Type 1 Diabetes association. (A) Pairwise scatter plots for proteins with real-data correlation ranging from [0.72, 0.8]. Except for streaks at 0, a Gaussian copula appears to preserve associations between pairs of proteins. (B) A heatmap of the correlation matrices across all proteins in the real and simulated data. Blocks of positively and negatively correlated proteins appear to be preserved in the simulation. (C) Prediction accuracy of SPLS-DA across simulation settings. The *x* and *y* axes correspond to sample size and AUROC, respectively. Color corresponds to the number of null features, 427 *×* (1 *− π*_1_), which governs the rate of power increase as a function of sample size.

We next altered the fraction denoted *π*_1_ of nonnull proteins whose means and standard deviations depend on T1D status. We chose the nonnull proteins according to their *p*-values from a Wilcoxon Rank-Sum test on the original experimental data. We considered the most significant proteins as true signals in the simulator. We then applied SPLS-DA with the pre-specified hyperparameters to semisynthetic data of varying sample sizes, with 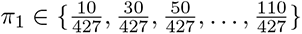, giving coverage of settings where differentially abundant taxa range from rare (*<* 2.5% of taxa) to relatively common (*>* 25% of taxa).

#### Data analysis results

When considering 20 samples total, we found that the proportion *π*_1_ of true nonnull proteins has little effect on average AUROC, which is only slightly better than random (Figure 4C). Larger *π*_1_ was associated with less variable performance. When 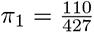, performance increased quickly with sample size, plateauing at 110 samples. In contrast, for 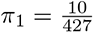, performance increased more gradually, with room for improvement even at 200 samples. Interestingly, performance was relatively stable from 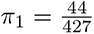 to 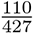. Altogether, this analysis suggested that if more than a few dozen proteins are thought to be associated with the outcome, then between 110 and 155 samples is sufficient. For fewer proteins, either a larger sample size or an alternative analysis strategy should be considered. A sense of the true proportion *π*_1_ of nonnull proteins can be gauged from the distribution of *p*-values in the template data, using estimators like those introduced in [99, 3, 4]. For these data, the true signal appeared to be weak. For example, Storey’s estimator returned 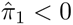.

#### Summary

We showed how semisynthetic data can help determine sample size for SPLS-DA applied to microbial proteomics. While no formulas for statistical power exist for this method, simulation allowed us to study how both experimental (sample size) and biological (proportion of nonnull proteins) factors influence its performance. Further, we detailed the process of evaluating the simulator’s correlation structure, demonstrating how visualizations and metrics could be used to iteratively refine a model.

### 4.3 Benchmarking network inference

#### Motivation

Network models give a holistic view of interactions in microbial ecosystems. They can identify tightly connected subcommunities and keystone taxa [41, 22, 77]. Unfortunately, validating these models is difficult, since determining ground-truth edges typically requires low-throughput experimentation such as knockout studies [9]. This creates a barrier to benchmarking, both for their use in specific studies and for evaluating new methodologies. Simulation can address these issues by providing ground-truth edges.

We compared two methods: SpiecEasi [55], a graphical lasso-based algorithm tailored to compositional data, and the Ledoit-Wolf estimator [60], a covariance estimator created for high-dimensional but low-rank data. A priori, we may expect SpiecEasi to perform better on microbiome data, since it was specifically designed for this purpose. However, this comparison has not been previously reported, and it is also unclear if potentially improved accuracy justifies the additional time required to solve the SpiecEasi optimization problem. To help answer this question, we can use simulations with known covariance structures.

#### Data

We analyzed the data from the American Gut Microbiome (AGM) project, a citizen science initiative where participants submit stool samples and complete detailed diet and health surveys [70]. The study has revealed associations between survey responses and microbiome composition. We considered a subset of 261 samples available through the SpiecEasi package [54], after excluding samples with fewer than 1000 reads. We filtered down to a “core” gut microbiome [93] of the 45 taxa with an abundance of over 100 in at least 50 samples.

#### Simulation and evaluation

We fitted a zero-inflated negative binomial GAMLSS model to these data using BMI and log sequencing depth as covariates. The BMI term was included because the original AGM study [70] found a significant association between BMI and microbiome composition. The sequencing depth term allows samples with deeper sequencing to have larger means on average across all taxa. The evaluations comparing real and simulated data are available in section 3.2 of the accompanying online tutorial. Here, we focus on benchmarking estimator accuracy across several network structures beyond those observed in AGM. We define several ground-truth correlation structures representing different statistical regimes: block, banded, Duncan-Watts small-world, scale-free, and Erdőos-ényi random graph structures [80]. The block and banded covariances were defined directly, while others were derived from Gaussian graphical models with the respective structure, shown in Figure 5A. For example, the scale-free network had several hub taxa with high covariance across many neighbors, while the Erdőos-ényi network connected all pairs of taxa with equal probability. Note that while our focus here is on covariance matrix estimation, a similar study using known, structured precision matrices could also be implemented.

**Figure 5:**
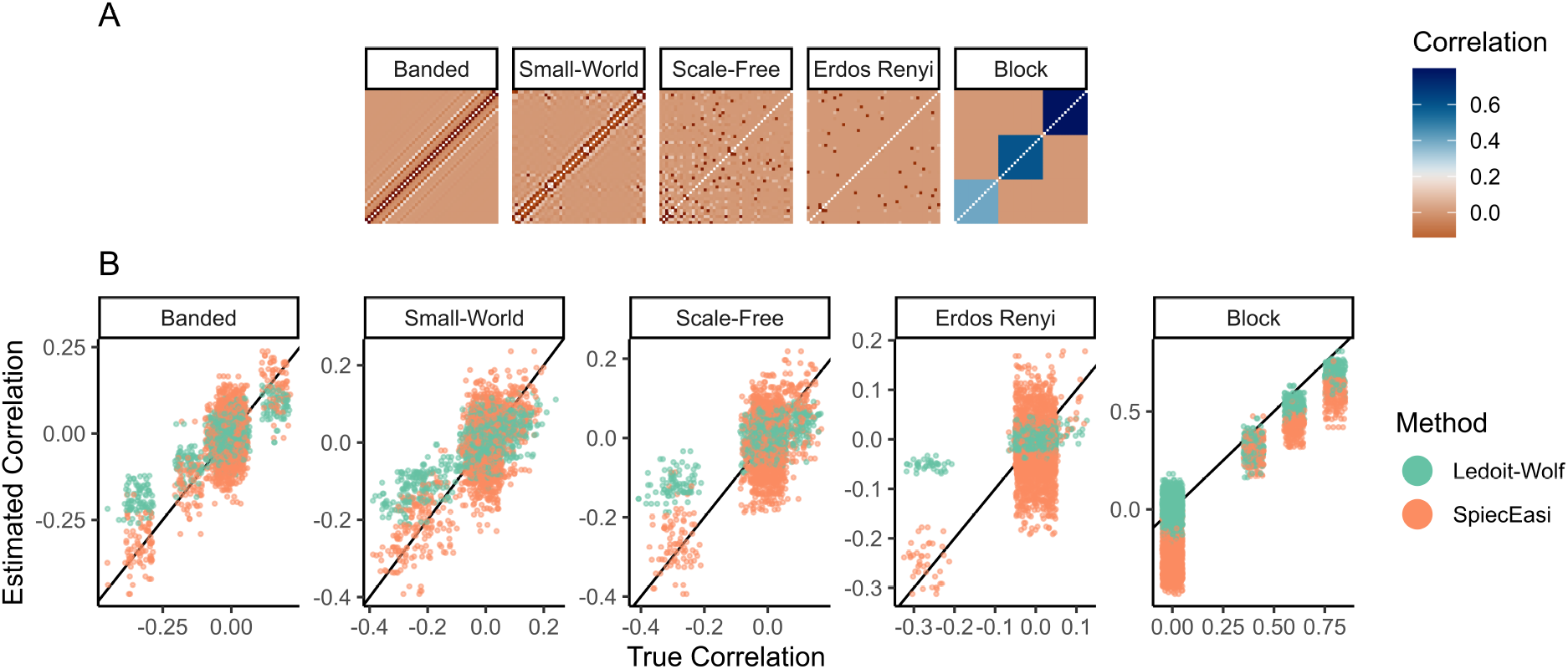
Benchmarking network methods for American Gut Microbiome project. (A) Ground truth correlation matrices used in the simulation study and (B) estimates made using SpiecEasi and the Ledoit-Wolf estimator. Entries (*j, j^′^*) in the matrices of (A) provide the copula correlation between taxa *j* and *j^′^*before transformation using the zero-inflated negative binomial model. From (B), SpiecEasi tends to have lower bias, since it is generally centered along the ground truth black line, but higher variance, since its vertical spread is larger. This figure also highlights settings where estimation tends to be easier (block and banded) and more difficult (Erdőos-ényi).

#### Data analysis results

Across network regimes, the Ledoit-Wolf estimator had larger bias but lower variance compared to SpiecEasi (Figure 5B). Since both estimators are regularized, they exhibited some bias towards correlations with smaller magnitudes. The Ledoit-Wolf estimator performed better in the banded and block settings, while the two approaches were comparable in other cases. All methods were challenged in the Erdőos-ényi setting, often declaring large correlations for taxa with no true relationship. Interestingly, in the block covariance scenario, SpiecEasi estimated many exact zero correlations as slightly negative. This is a consequence of SpiecEasi’s compositional assumption, which induces negative correlation across taxa. In spite of this artifact, the block covariance setting appeared to support more efficient estimation than any of the other covariance settings we considered. Thus, if we assume that a community has a block correlation structure, then fewer samples may be required. This is consistent with general statistical theory, which argues that low-dimensional block structure can greatly simplify high-dimensional covariance estimation [84].

#### Summary

We showed that simulations offer a useful lens for comparing network inference methods across diverse network structures while maintaining realistic abundance distributions. Starting from a single template dataset, we were able to simulate according to several types of ground truth correlation, enabling us to identify settings where methods are more likely to fail or succeed.

### 4.4 Batch effect correction

#### Motivation

In large microbiome studies, it is often difficult to guarantee uniform data collection and processing for all samples, leading to systematic differences across experimental groups, often referred to as batch effects [103, 66]. For example, storing samples at different temperatures might change the abundances of taxa whose marker gene sequences degrade more rapidly at some temperatures. Failing to address these batch effects can limit power by masking treatment effects. They can also compromise validity by introducing spurious differences across treatment groups [61]. Hence, a common preprocessing step in microbiome data analysis is to apply batch effect correction to standardize data across batches [30, 64].

Despite their increasingly widespread adoption in microbiome studies, batch effect correction methods must be applied carefully. Failing to account for the correction in downstream differential tests can lead to miscalibrated *p*-values [75] or performance estimates [96]. Further, it can be difficult to balance underintegration, where batch effects persist post-correction, against overintegration, where aggressive batch effect correction eliminates meaningful biological variation [113]. It is often unclear in advance whether these issues will arise for a given dataset or batch effect correction method. By defining ground-truth batch and biological effects, simulations give a way to evaluate batch effect correction methods and their impact on downstream inferences. Moreover, they allow quick comparison of methods across experimental designs and biological scenarios, allowing more complete evaluation than is possible in isolated benchmark datasets.

#### Data

We analyzed a study of anaerobic digestion (AD). AD is a biodegradation process underlying many bioenergy production technologies. The study Chapleur et al. [6] profiled the microbiomes from AD samples treated with phenol, a micropollutant that affects biodegradation efficiency and stability, posing challenges for large-scale industrial deployment. We use a subset of 75 samples and 231 genera discussed in Wang and Lê Cao [103], which focused on changes in community composition under two phenol concentrations. Since obtaining samples is time-consuming, the experiment was carried out over five sessions, leading to batch effects visible in the PCA plot in Figure 6A.

**Figure 6:**
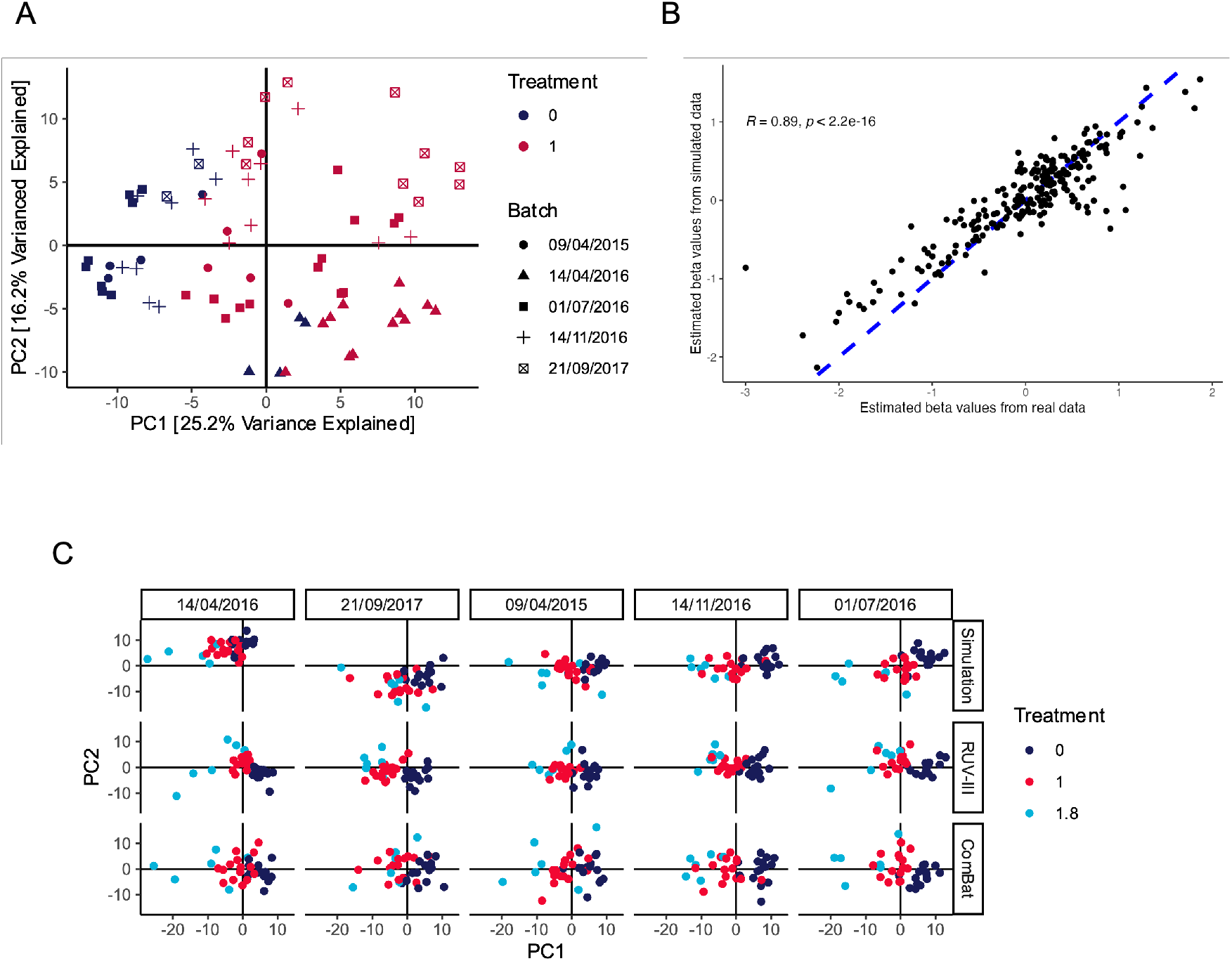
Assessing batch effect correction methods in an anaerobic digestion study. (A) A principal components plot of the original AD dataset reveals significant batch-to-batch variation. For example, all experimental samples generated on 14/04/2016 are shifted towards the bottom-right quadrant. Note that the technical replication structure is also visible, for example, the clusters of untreated samples from 01/07/2016 on the left-hand side. Though the two phenol concentration treatments can still be distinguished from one another, removing the batch effect can improve power. (B) Narrow evaluation using a scatterplot and correlation coefficient to compare the RUV-III batch effect regression coefficients for each taxon in real and simulated data. (C) Original and batch corrected data in the simulation with a new, less frequently sampled treatment group (*t* = 1.8) introduced. Batches (columns) have been sorted from the lowest to the largest average PC1. Both RUV-III and ComBat successfully remove systematic differences between batches, but in some cases ComBat removes true differences between the *t* = 1 and *t* = 1.8 groups. Note that the principal components are derived separately for each row.

#### Simulation and evaluation

The purpose of this simulation experiment is to compare the performance of RUV-III [73] and ComBat [47], two batch-effect correction methods, when applied to data like those in the AD study. We used the CLR-transformed AD dataset as our simulation template. For each taxon, we applied a Gaussian GAMLSS and copula model with batch and treatment status as covariates. Given the relatively small sample size, we used an adaptive thresholding covariance matrix estimator [5] within the Gaussian copula. PCA on data simulated from this model revealed that the simulator could recapitulate the observed batch effects. We can evaluate this more precisely using a narrow utility evaluation, comparing the RUV-III batch effect regression coefficients from the real and simulated data. For each taxon, a regression coefficient was calculated using AD as the factor of interest. The regression coefficients from the simulated data closely matched those from the real data, with an overall correlation of 0.89 (Figure 6B).

Batch effect correction methods are known to be sensitive to imbalance between true biological groups [66]. We next explored whether this could pose a challenge in AD studies if we considered an additional phenol concentration treatment level. We simulated a hypothetical scenario where a stronger treatment had been applied to a small subset of samples. Each batch was assigned 15 samples at the reference concentration (*t* = 0) and the previously observed comparison group (*t* = 1), but only six at a new, hypothetical treatment (set to *t* = 1.8). Relative to the reference, this new treatment level is anticipated to perturb the microbiome community in the same direction as the original *t* = 1 group, but to a greater extent. Since this treatment group is less frequently sampled, there is a risk that its effect might be masked by overly aggressive batch integration.

#### Data analysis results

We compared the results of RUV-III and ComBat applied to data simulated from our altered experimental design (Figure 6C). The simulator generated data with clear batch and treatment effects (first row). Although we observed a lack of replication structure for some of the batches in Figure 6A, the simulated data preserved sufficient batch effects to evaluate batch effect correction methods. We then compared the PCA projections when RUV-III and ComBat were applied to the simulated data. Both methods centered all batches around the origin, as expected from a batch effect correction method. However, the *t* = 1.8 group was less clearly separated from the *t* = 1 group in the ComBat output compared to either the original simulation or the RUV-III corrected data. For a more quantitative analysis, we trained a linear discriminant analysis to predict the treatment group from the first two principal components, yielding a classification accuracy of 73.9% on the original simulation, 78.3% on the ComBat correction, and 88.3% on the RUV-III correction. These classifications performance results suggest that ComBat may overintegrate imbalanced data, while RUV-III preserves more subtle treatment differences.

#### Summary

We illustrated how simulation can give insight into the behavior of candidate batch integration methods in a way that is tailored to specific data analysis contexts, with the opportunity to create new treatment scenarios not present in existing real-data benchmarks.

### 4.5 Omics data integration

#### Motivation

A single assay can only give a partial view of a microbial ecosystem. For example, 16S rRNA sequencing data characterizes bacterial community composition within a sample, but other kingdoms, metabolites, and host cells shape microbiome properties. To capture these features, different sequencing methods conducted on the same samples (e.g., ITS for fungi) or profiling techniques (e.g., NMR for metabolites) are needed. By analyzing the complementary views offered from diverse datasets, it is possible to develop a more holistic understanding of an ecosystem’s biology [90, 102, 8].

Effectively analyzing these data, however, is challenging, with no consensus on how integration strategies should be deployed across contexts. The statistical task is not simply multivariate but also multiassay, linking tables influenced by diverse biological or experimental factors. It is necessary to decide on which datasets to integrate, how each table should be normalized, and which integration method might be appropriate. Simulation can give a controlled, simplified setting within which to compare analysis strategies and can help to predict method performance under realistic biological scenarios.

Integration methods often return a dimensionality reduction plot designed to uncover shared covariation across assays, analogous to how PCA arranges samples to describe variation within a single assay. It is critical that these methods faithfully preserve between-assay and between-sample relationships. Ideally, data integration dimensionality reduction will highlight genuine, shared axes of variation across data sources while avoiding the appearance of false associations between unrelated data sources. We design a simulation to assess the extent to which these plots can misleadingly suggest similarities in inherently “unalignable” assays [67]. This multi-assay analysis parallels the multi-batch problem discussed in the previous case study – instead of studying overintegration across batches, we consider overintegration across assays.

#### Data

We re-analyzed the data from Haak et al. [36], who studied how the gut microbiome is altered during sepsis in ICU patients. Sepsis can be triggered by microbial infection unrelated to bacteria (e.g., from the *Candida* fungus). Therefore, each sample was profiled using ITS amplicon and Virome sequencing in addition to 16S rRNA sequencing. Since sepsis is treated with antibiotics, the study included healthy patients undergoing antibiotics treatment. The study included 20 healthy controls, 23 sepsis patients, 5 healthy patients on antibiotics, and 9 ICU patients without sepsis. After applying the same filtering as [36], we obtained measurements for 180 bacterial genera, 18 fungal genera, and 42 viruses.

#### Simulation and evaluation

We considered the multiblock SPLS-DA method to integrate these class-labeled sequencing datasets [94], and simulated from the scenario where the class label was relevant only for a subset of tables. This could occur if any of the three kingdoms assayed were unrelated to sepsis or antibiotics. An ideal multiassay integration in this setting would recover the similarities in the abundance profiles across kingdoms (e.g., revealing shared clusters of samples) while avoiding introduction of false relationships with sepsis or antibiotic status. Note that multiblock SPLS-DA is the multi-assay extension of SPLS-DA (discussed in the section ‘Power analysis for multivariate methods’) and replaces the SPLS-DA objective with a weighted average of covariances across pairs of tables. We applied a Gaussian GAMLSS simulator to each assay, as they were already normalized. For the ITS and Virome assays, we allowed means and variances for each feature to depend on a categorical feature encoding both ICU and antibiotic use. For the 16S data, we used two simulators: one which conditioned taxonomic abundances on ICU/antibiotics category and one which did not. We joined all tables using a Gaussian copula with an adaptive covariance estimator to handle high-dimensionality. Therefore, in the ground truth simulation, the marginal (taxon-level) and covariance (community-level) structure from the template data are maintained. However, while the sepsis and antibiotic factors continue to influence all the ITS and Virome measurements, this relationship has been deliberately removed from one of the two simulated 16S datasets.

This design allows us to study the extent to which integration can spuriously introduce sepsis or antibiotic associations into the 16S data. Specifically, we then compared multiblock SPLS-DA output on the original and both versions of the simulated data (Figure 7).

**Figure 7:**
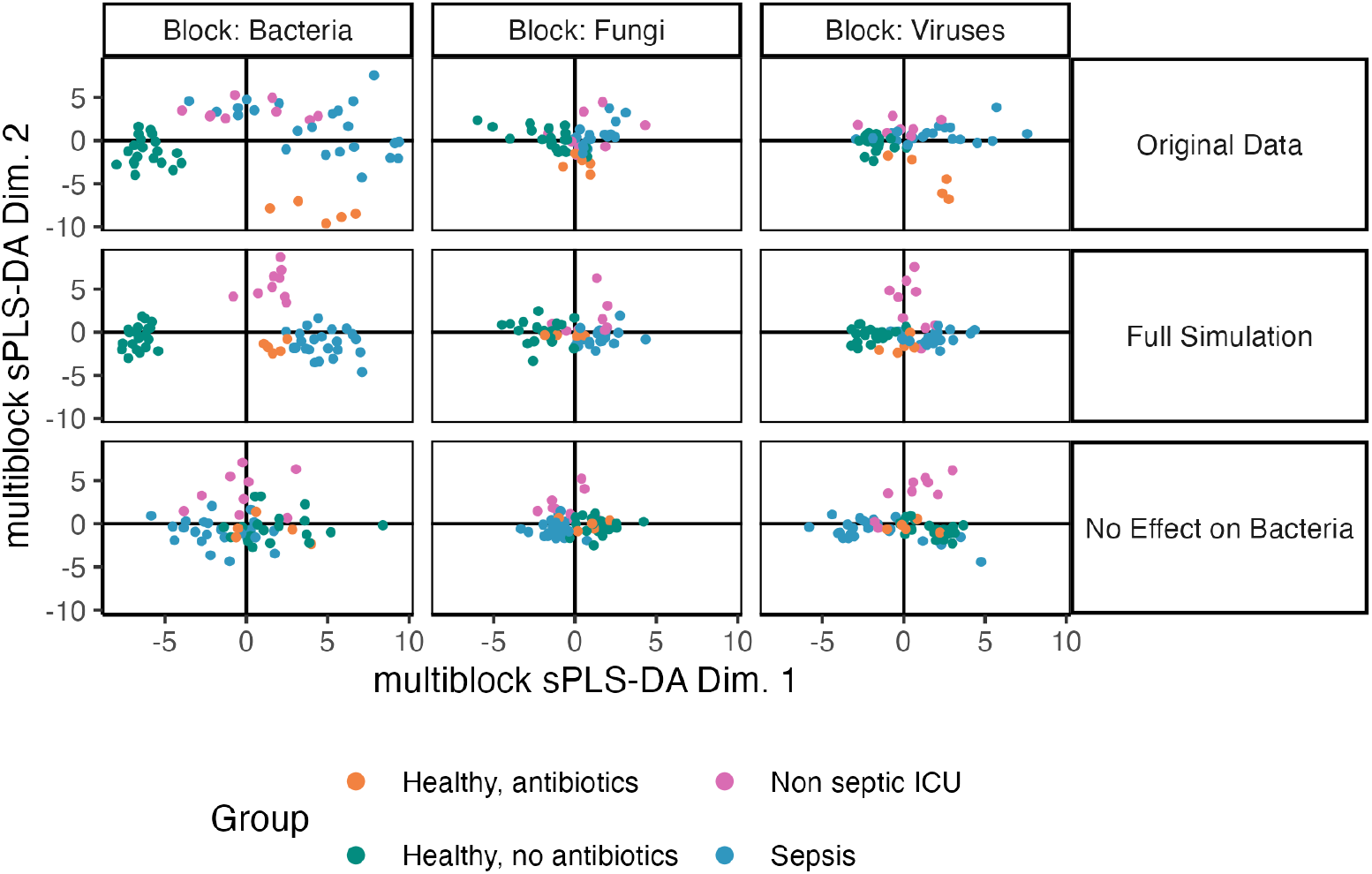
Using simulation to understand properties of omics data integration methods in a transkingdom analysis. Columns correspond to three amplicon sequencing datasets for transkingdom analysis of sepsis from Haak et al. [36]. Multiblock SPLS-DA projections from real and simulated data are given in the first two rows. The bottom rows show a perturbed simulation where all associations in the 16S data have been removed. The fact that between-group differences are still visible reflects the cross-table reduction and serves as a null reference for reliability analysis.

#### Data analysis results

We find that, even after removing all association between 16S abundances and the class label, the multiblock SPLS-DA projections for the 16S block still separated by class, albeit more weakly than before. This is a consequence of the analysis’ multiblock nature: strong class associations from other tables can be artificially introduced into the 16S table. Indeed, part of the objective of the multiblock SPLS-DA algorithm is to maximize similarity in projections across all tables. In this case, in the real data, the class differences in the 16S data were much larger than those seen in this null scenario. Thus, simulation can guarantee reliable conclusions for complex integration tasks.

Further, to decide on which tables should be analyzed together, or separately, we calculated alignment measures [67] on the real data and compared them with a synthetic null where the tables are not alignable. We study the distribution of canonical correlations in both the observed and synthetic negative control data when integrating the Virome and 16S data together (Figure 8). The control was defined so that the two tables are known to have no correlation within the underlying copula model. The results showed the presence of slight, but systematic, shared variation: observed canonical correlations were consistently larger than those in the synthetic null for the top dimensions, but the increase was quite modest. This suggests that substantial variation is isolated within the tables in a way that an integrated analysis cannot capture, motivating the use of within-table analyses alongside integrative methods.

**Figure 8:**
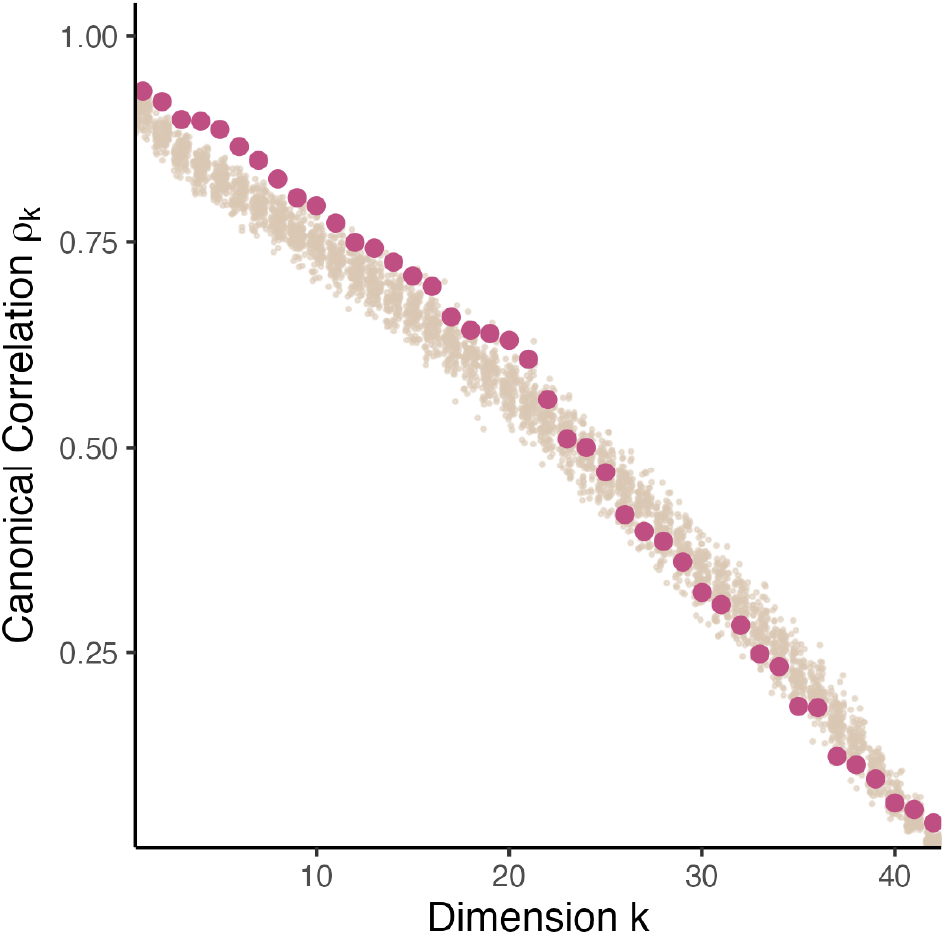
Comparison between canonical correlations in real (red) and reference null (beige) samples in a transkingdom analysis. For the first twenty dimensions, the real canonical correlations are slightly larger than those in the synthetic null. These dimensions are likely to reflect true, if modest, shared variation across tables.

#### Summary

Analogous to using negative controls in biological experimentation, we created synthetic negative controls in a computational workflow for data integration. This allowed us to gauge evidence for potential discoveries in real data, and prioritize either simultaneous or table-specific analysis based on departures from the synthetic negative control setting.

## 5 Discussion

We have reviewed the potential to apply simulators to microbiome study design and analysis. We have considered advances that make simulators more broadly applicable than their predecessors and have provided reproducible case studies showing how they can offer ground truth for evaluating FDR and power (sections ‘Benchmarking differential abundance methods and statistical power for univariate methods’ and ‘Power analysis for multivariate methods’), help identify regimes where competing methods differ (sections ‘Benchmarking network inference’ and ‘Batch effect correction’), and ensure valid interpretation of statistical outputs (section ‘Omics data integration’).

We have emphasized methods for evaluating simulation quality, introducing vocabulary for distinguishing between fit-for-purpose, narrow, and global metrics. These metrics can be used both to compare simulation packages, like those listed in Table 2, and to refine initial simulator formulations. Moreover, this review has identified criteria for determining a simulator’s relevance to specific applications, including distributional assumptions, ability to match a template, handling of multivariate relationships, and incorporation of experimental or biological effects. Between simulators, there are trade-offs between faithfulness, controllability, and generality of application, and the most appropriate approach will depend on the relative importance of these factors.

A key insight of modern simulation is that models can be trained on real experimental data from related contexts. A semisynthetic approach can enhance simulation quality and requires less implementation effort compared to designing a de novo mechanism. However, it is worth cautioning that even a simulator that perfectly emulates a template dataset may still be unhelpful if the template is not a good match to the motivating problem. There is a trade-off between using template data from pilot experiments, which are small but highly representative of future conditions, and large public databases, which offer more data but may not directly relate to the problem of interest.

We expect future simulators to support workflows with novel data modalities and their combinations.

For example, while we considered integration across batches and assays, their heterogeneity was mild. Developing simulators that model variation not just across batches but across cohorts, and not just for different amplicon technologies but for entirely different assay types, is a worthwhile direction for further study. Moreover, though we focused on simulating community taxonomic and metabolomic profiles, methods for simulating metagenomics reads have also been proposed [40, 46, 34], and incorporating template data into read simulation could allow systematic study of entire processing and analysis pipelines, similar to recent advances in single-cell read simulation [107].

As simulators become easier to develop and apply, the potential for reusing others’ work will increase. For example, instead of creating one-off simulators for the power analysis in a grant proposal or the simulation study in a methods paper, researchers could borrow and modify existing simulator definitions and output. Curating repositories of reusable simulators, where the template data and generating mechanisms are specified, would be valuable. While we have focused on researcher-level power analysis, benchmarking, and reliability analysis, simulation also has the potential to resolve field-level controversies. For example, no consensus has emerged for the analysis of strain-level variation, with some researchers emphasizing their importance to health outcomes [108] and others cautioning against attempts to detect them with existing technology [45, 53]. Simulation can provide the ground truth and control necessary for fine-grained discussion of these issues. Indeed, it has already played an important role in a debate about the use of supervised normalization in microbiome data [101, 13]. As techniques become more powerful and accessible, computational studies using semisynthetic data will become an important part of microbiome research toolkit.

## Notes

https://krisrs1128.github.io/microbiome-simulation/

https://github.com/krisrs1128/microbiome-simulation/

### Competing Interest Statement

The authors have declared no competing interest.

